# Seasonal plasticity in neuroendocrine mechanisms relevant to year-round territorial aggression in a wild teleost fish

**DOI:** 10.1101/2025.10.23.684146

**Authors:** G. Valiño, C. Jalabert, J. Farías, J.R. Sotelo-Silveira, K.K. Soma, L. Quintana

**Affiliations:** Departamento de Neurofisiología Celular y Molecular, Instituto de Investigaciones Biológicas Clemente Estable, MEC, Montevideo, Uruguay; Grupo Cronobiología, CSIC, Facultad de Ciencias, Universidad de la República, Montevideo, Uruguay; Espacio de Biología Vegetal del Noreste, CENUR Noreste, Universidad de la República, Tacuarembó, Uruguay; Departamento de Genómica, Instituto de Investigaciones Biológicas Clemente Estable, Montevideo, Uruguay; Department of Psychology, University of British Columbia, Vancouver, BC, Canada

## Abstract

Animals experience cyclical environmental changes, such as seasons, that require physiological adjustments to support different behaviors. Although many behaviors occur only during specific periods, some species, like *Gymnotus omarorum*, display territorial aggression year-round, making them valuable models to study seasonal plasticity in the mechanisms maintaining stable behavioral outputs. *G. omarorum* is a teleost fish in which neuroestrogens have been shown to play a key role in non-breeding aggression. Here, we quantified circulating hormone levels and gene expression in the social behavior network of wild breeding and non-breeding individuals. During the non-breeding season, both sexes exhibited elevated circulating androgen levels, providing potential substrates for local estrogen synthesis. Consistently, brain aromatase and estrogen receptor expression were also upregulated, suggesting an increased capacity for local estrogen synthesis and signaling. Our findings provide the first evidence in a teleost of seasonal plasticity in the mechanisms underlying territorial aggression. Comparisons with birds and mammals reveal both shared and lineage-specific strategies, highlighting common endocrine principles while revealing the evolutionary diversity of solutions to maintain a stable behavioral phenotype across changing seasonal contexts.

## INTRODUCTION

Seasonal plasticity, also known as life-cycle staging, encompasses cyclical changes in physiology, morphology, or behavior that occur at predictable times of the year [1,2]. This form of phenotypic flexibility allows individuals to adjust to cyclical environmental challenges such as seasonal shifts in temperature, photoperiod, and resource availability. These adjustments often manifest as pronounced seasonal variations in social behaviors such as reproduction, migration, or parental care. Accordingly, the physiological mechanisms that sustain these behaviors are also expected to vary seasonally, including changes in key neural circuits such as the social brain network (SBN, [3,4]), often mediated by sex steroids [5]. However, some animals express certain social behaviors year-round, such as territorial aggression, even as circulating sex steroid levels fluctuate seasonally [6]. This observation challenges the prevailing assumption that seasonal social behaviors depend exclusively on circulating hormonal control and raises the possibility of alternative neural or physiological mechanisms. When a behavior remains consistent across life-history stages, the underlying mechanisms supporting it may differ [7], suggesting that territorial behavior is likely regulated by distinct seasonal endocrine mechanisms despite showing similar behavioral traits across seasons.

During the breeding season, circulating androgens and estrogens facilitate territorial aggression in both males [8–10] and females [11–13]. In contrast, non-breeding territorial aggression occurs when gonads are regressed and circulating sex hormones are low or undetectable, and even persists after castration, indicating that gonadal steroids are not required [14–16]. However, pharmacological manipulations have shown that non-breeding aggression depends on estrogens and androgens across birds, mammals, and fish [16–20], indicating that non-breeding territorial aggression is regulated by non-gonadal sex steroids.

The brain is known to be a site of sex steroid synthesis [21–23]. The capacity for local steroid synthesis is deeply conserved across vertebrate taxa, with key steroidogenic enzymes expressed throughout the brains of fish, amphibians, birds, and mammals [24]. Although this steroidogenic machinery is present year-round, evidence suggests it plays a critical regulatory role in maintaining non-breeding territorial aggression [25,26]. This has led to the proposal of a seasonal alternation in the mechanisms that maintain year-round territorial aggression [27–29]. Supporting this mechanistic shift, several non-exclusive changes have been documented. Peripheral tissues can increase the production of weak androgen precursors during the non-breeding season [30–34] that are converted within the brain into active steroids via local steroidogenic enzymes. The brain’s capacity for local steroid synthesis can itself be upregulated seasonally [25,26,35]. Additionally, neural sensitivity to these hormones can increase through upregulation of estrogen and androgen receptors [36–40]. Collectively, these mechanisms reveal remarkable neuroendocrine plasticity in the regulation of year-round territorial aggression. However, little is known about this phenomenon outside birds and mammals, limiting our understanding of how widespread these neuroendocrine strategies are across vertebrates.

The mechanisms underlying seasonal changes in territorial aggression remain poorly understood in teleost fish, which represent the largest and most diverse group of vertebrates. *Gymnotus omarorum* is a weakly electric fish from South America in which both males and females exhibit year-round territorial aggression. *G. omarorum* is a well-established teleost model for studying year-round territorial aggression, integrating behavioral, neuroendocrine, and ecological approaches across field and laboratory contexts [41–45]. Field studies show that males and females maintain spatial distributions consistent with year-round territoriality [46], corroborated by individual tracking data [47]. In laboratory settings, both sexes display robust aggression outside breeding contexts, thoroughly characterized in terms of locomotor displays and electrical social signaling [48–50]. This aggression remains unaffected by castration [16]; nevertheless, pharmacological manipulation of both androgenic and estrogenic pathways modulates this behavior in males and females alike [16,19,20]. Consistent with a role for neural steroid synthesis, both sexes exhibit elevated forebrain estrogen concentrations during the non-breeding season despite undetectable circulating levels [51]. Altogether, these features make *G. omarorum* an exceptional model for investigating the neuroendocrine mechanisms that sustain stable behaviors across changing seasonal contexts.

Here, to test whether a seasonal shift in hormonal substrates underlies the persistence of territorial aggression across reproductive contexts, we quantified circulating sex steroids and measured the expression of aromatase and sex steroid receptors in specific nuclei within the SBN in wild-caught males and females during the breeding and non-breeding seasons. We hypothesized that there would be an increase in circulating precursors and in the gene expression of aromatase and/or sex steroid receptors in the SBN of both males and females during the non-breeding season, in comparison to breeding levels.

## MATERIALS AND METHODS

### Animal collection and sampling

Wild adult females and males were captured during the breeding (4 January 2019) and non-breeding season (18 July 2019) at Laguna de los Cisnes, Maldonado, Uruguay (34°48’S, 55°18’W). Sample sizes were: non-breeding females (n = 11), breeding females (n = 15), non-breeding males (n = 13), and breeding males (n = 12). Fish were collected during the daytime, which corresponds to their resting period [47,52]. Individuals were located without disturbance using a detector that converts electric organ discharges into audible signals and were then captured individually with a rigid hand net, following [53].

### Sample processing and preservation

Blood and brain extractions followed the procedures described by [51]. Briefly, once fish were netted, they were rapidly anesthetized by immersion in eugenol solution (1.2 mg/L). Blood was drawn from the caudal vein with a heparinized syringe and kept on wet ice. Fish were then decapitated, and brains were rapidly removed, placed in powdered dry ice for flash-freezing. To minimize handling-induced stress, blood was extracted and animals decapitated within 3 min of capture, and brain dissection was completed within 5 min of capture. Blood samples were centrifuged (14,000g for 10 min) and plasma was obtained. Once in the lab, plasma and brain samples were stored at –80°C. Gonads were also dissected for sex confirmation and gonadosomatic index calculation ([gonad weight / body weight] × 100).

All procedures followed ASAP/ABS Guidelines for the Use of Animals in Research and were approved by the Institutional Ethical Committee (Comisión de Ética en el Uso de Animales, Instituto Clemente Estable, MEC, CEUA-IIBCE: 001/02/2018).

### Steroid hormone extraction and quantification

A panel of eight steroids was quantified: androstenedione (A4), testosterone (T), 11-ketotestosterone (11-KT), dehydroepiandrosterone (DHEA), estrone (E1), 17β-estradiol (E2), progesterone and cortisol. Steroids were extracted from 25 μL of plasma using liquid-liquid extraction followed by solid-phase extraction, as previously validated in this species (Zubizarreta & Jalabert et al., 2023). Briefly, samples were placed in polypropylene vials containing zirconium ceramic beads, and deuterated internal standards (progesterone-d9, cortisol-d4, DHEA-d6, testosterone-d5, estradiol-d4) in 50% MeOH were added to correct for analyte losses and matrix effects. Testosterone-d5 served as the internal standard for A4, T and 11-KT; estradiol-d4 for E1 and E2; and progesterone-d9, cortisol-d4, and DHEA-d6 for their respective analytes. Steroids were extracted twice with ethyl acetate to increase extraction efficiency, washed with water, and dried under vacuum at 60°C. Dried residues were reconstituted in methanol and subjected to C18 solid phase extraction. Samples were eluted with methanol, dried under vacuum, reconstituted in 25% methanol, and centrifuged. The final supernatant was transferred to LC vial inserts and stored at –20°C until analysis. Plasma samples were processed alongside standards, quality controls, blanks, and double blanks. The coefficient of variation between quality control replicates for all steroids was <14%.

Steroids were quantified on a Sciex 6500 Qtrap triple quadrupole tandem mass spectrometer, as previously described for this species [51]. Briefly, 45 μL of each extract were injected into a UHPLC system and separated on a C18 column using a methanol/ammonium fluoride water gradient as a modifier. Two multiple reaction monitoring transitions were used for each steroid and one for each deuterated internal standard. Data were acquired in positive electrospray ionization mode for all steroids except E1 and E2, which were analyzed in negative mode. The lower limit of quantification was 0.008 ng/mL for cortisol, T, 11-KT, and E1; 0.016 ng/mL for progesterone, A4 and E2; and 0.16 ng/mL for DHEA. All blanks and double blanks were below the lowest standard on the calibration curves.

### Brain microdissection

Brains were sectioned in the coronal plane at a thickness of 200 µm using a cryostat (SLEE Mainz, MEV), and sections were collected onto glass slides previously cleaned with 70% ethanol and sterilized with UV light. Three regions of the social brain network were selected for microdissection based on [54]: the ventral nucleus of the ventral telencephalon (Vv; homologous to the lateral septum in mammals and birds), the ventral supracommissural nucleus (Vs; homologous to the bed nucleus of the stria terminalis in mammals and birds), and the preoptic area (POA). Nuclei were isolated using the Palkovits punch technique (300 µm, inner diameter of punch tool; [55]). Tissue was collected into 1.5 ul tubes and stored at −80°C until mRNA extraction.

### Gene expression quantification

Primer pairs for qRT-PCR were designed from the *G. omarorum* transcriptome [56] using Primer3. The primers employed were *β-actin* (F: 5′-GTGCCCATCTACGAGGGTTA-3′; R: 5′-GCTGTGGTGGTGAAGCTGTA-3′), *cyp19a1b* (F: 5′-TGGAGAGCTGTCAGTGGATG-3′; R: 5′-CTGTTTCAGGAGCACCAACA-3′), *esr1* (F: 5′-GAGGAGAAGCTGTGCCTGTC-3′; R: 5′-CTTCATGGTGCAGTTCATGC-3′), *esr2b* (F: 5′-CCTGACCTTCTACAGCACACC-3′; R: 5′-ACTATACACCAGCGGGGATG-3′), and *ar2* (F: 5′-AAAACAGCAAAACGGAGCAG-3′; R: 5′-ACCACTGTGTGCCACTTCTG-3′). Primer specificity was verified by sequencing the qRT-PCR amplicons (Macrogen). Amplification efficiencies were within the acceptable range (*β-actin*: 105%, *cyp19a1b*: 103%, *esr1*: 98%, *esr2b*: 100%, *ar2*: 100%). *β-actin* was used as the reference gene for normalization as its expression did not differ among seasons or sexes across all three brain regions examined.

Total RNA was extracted with the RNAqueous™-Micro Total RNA Isolation Kit (Ambion, AM1931) according to the manufacturer’s protocol and reverse-transcribed into cDNA using the SuperScript™ III First-Strand Synthesis System (Thermo Fisher). qRT-PCR reactions were performed in CFX96 Deep Well^TM^ (Bio Rad), in duplicate in 20 µL volumes containing 10 µL of SYBR™ Green Universal Master Mix (Applied Biosystems™), 0.8 µL of each primer (400 nM), 2 µL of cDNA (normalized at retro-transcription step), and 6.4 µL of Milli-Q water. Thermal cycling conditions consisted of an initial denaturation at 95 °C for 5 min, followed by 40 cycles of 95 °C for 15 s and 60 °C for 60 s. A melt curve was then generated by increasing the temperature from 65 °C to 95 °C in 0.5 °C increments. Relative gene expression was quantified using the 2^−ΔΔCt method [57]; the reference group varied among different comparisons and is stated in the figure legend of each qRT-PCR figure.

### Statistical analysis

For circulating hormones, values below the lowest standard on the calibration curve were considered below the lower limit of quantification (LLOQ). We quantified 8 hormones in 4 experimental groups. Of these 32 datasets, 9 presented 100% of detectable values, 4 presented at least 75% of detectable values, 1 presented at least 50%, 2 datasets presented at least 25% of detectable values, and 16 datasets presented undetectable values. In cases when ≥ 20% of samples in a dataset were above the LLOQ, then values below the LLOQ were estimated via quantile regression imputation of left-censored data using the MetImp web tool following [58,59]. When < 20% of samples in a group were above the LLOQ, then values below the LLOQ were set to 0.

To visualize patterns of circulating hormones and gene expression across experimental groups, radar plots were generated. Plasma hormone concentrations were log-transformed prior to analysis. For qRT-PCR data, relative expression was calculated as –ΔCt values. Median values were computed separately for each experimental group (non-breeding females, breeding females, non-breeding males, and breeding males). Median values for each group (non-breeding females, breeding females, non-breeding males, and breeding males) were first calculated and then standardized (z-scored) across all four groups for each variable, using the mean and standard deviation of the group medians. This standardization provided a common baseline for comparison, allowing hormonal and gene expression profiles to be visualized on an integrated scale (±3 SD) in the radar plots. To test the effects of season and sex upon the hormonal and gene expression profiles we performed permutational multivariate analysis of variance (PERMANOVA, [60]) using Euclidean distance with 999 permutations, in a four-group analysis (breeding females, non-breeding females, non-breeding males and breeding males).

For univariate comparisons of circulating hormone levels and gene expression, data normality was tested with the Shapiro–Wilk test. When both groups showed normal distributions, group differences were assessed using Welch’s *t*-test; otherwise, the Mann–Whitney *U* test was applied. When comparing groups in which one group had all samples below the LLOQ, Fisher’s exact test was applied. Statistical significance was set at p < 0.05 for PERMANOVA, Fisher’s exact test, Welch’s *t*-test and Mann–Whitney tests. To evaluate relationships between circulating hormones and brain gene expression, Spearman correlations were performed, with statistical significance considered at p-adjust < 0.05 following the Benjamini–Hochberg False Discovery Rate procedure. All statistical analyses and visualizations were performed in Python.

### AI assistance disclosure

Artificial intelligence tools (ChatGPT-4/5, Claude Sonnet 4, DeepSeek 3.1) were used to improve English grammar and assist in code development for statistical analyses and figure generation.

## RESULTS

### Global patterns of gene expression and hormones across groups

As a first approach to broadly visualize global patterns across circulating hormones and brain gene expression, we generated a radar plot for each group. Wild breeding and non-breeding females and males showed unique multi-hormonal and gene expression profiles (Fig. 1). To compare relative shapes and profiles, group median values were standardized (z-scored) across all four groups for each variable, allowing direct visualization of relative differences in hormonal and gene expression patterns. Notably, group profiles displayed similarities by season rather than by sex, indicating that season exerts a stronger influence on the global pattern. Both non-breeding females and non-breeding males showed a broad distribution with values above the reference for gene expression in Vs and Vv as well as for circulating androgens, with a notable absence in estrogenic hormones. While radar plots of breeding groups show a striking difference in shape with the non-breeding groups. Breeding females and males showed an overall very moderate gene expression in brain areas, while their circulating hormone profiles (although different) included components above average. This visual clustering pattern was statistically confirmed by PERMANOVA, which showed significant seasonal effects on hormonal and gene expression profiles (F = 5.85, p = 0.001) while sex differences were not significant (F = 1.19, p = 0.262).

**Figure 1:**
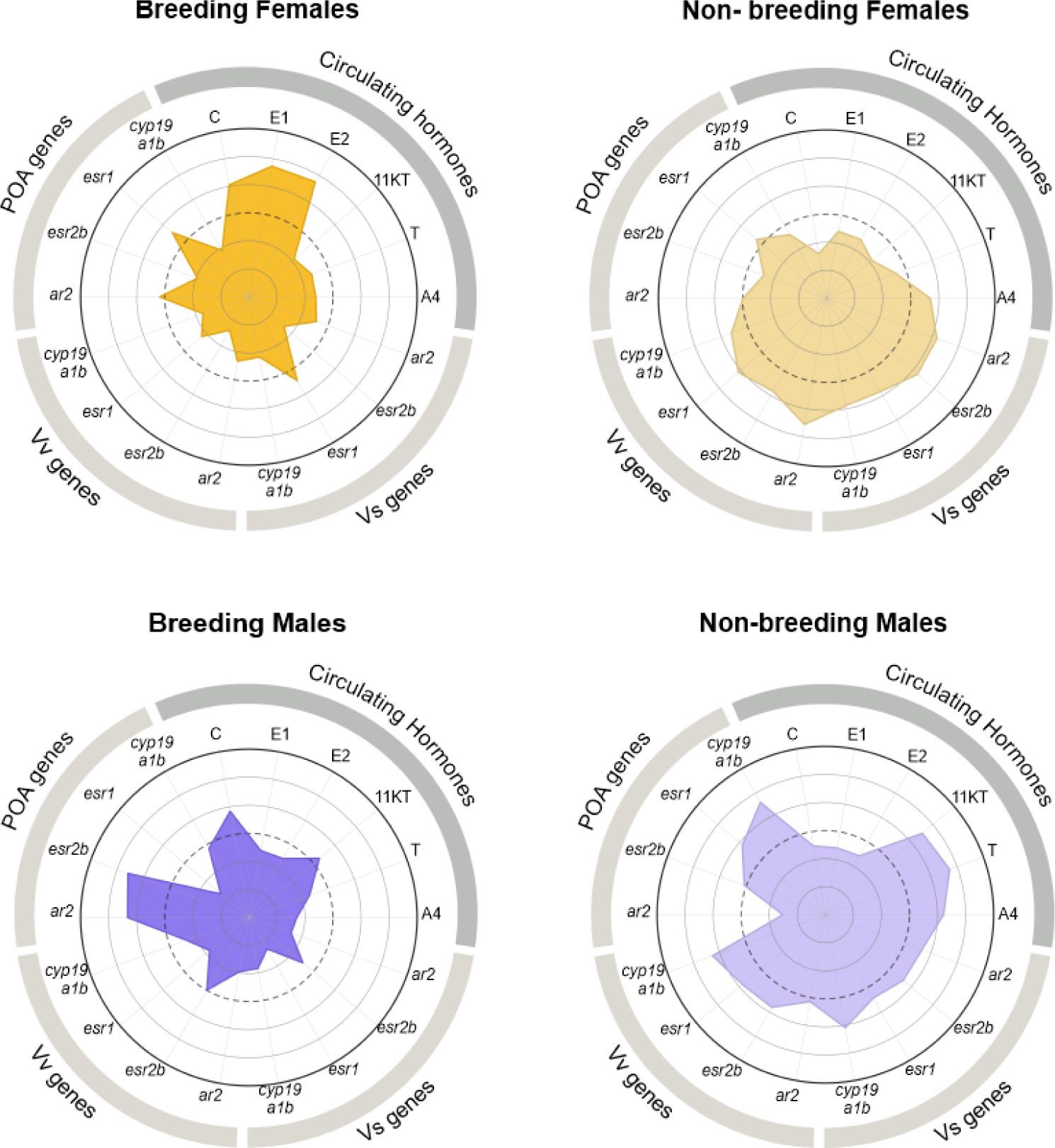
Radar plots depicting patterns across circulating hormones and brain gene expression for the four experimental groups. Each variable was normalized to the median value across all groups for visualization purposes. The dashed circular line represents this standardized group median (0). Concentric circles denote standardized levels in units of standard deviation (σ) from the group median, with values above 0 indicating higher relative levels and values below 0 indicating lower relative levels (circles from inner to outer: −2σ, −1σ, 0, +1σ, +2σ). Profile shapes are more similar within seasons than between sexes, suggesting seasonality has a stronger impact than sex on the overall expression pattern. Abbreviations: C: cortisol; E1: estrone; E2: 17-β estradiol; 11-KT: 11-ketotestosterone; T: testosterone; A4: androstenedione; *ar2*: androgen receptor 2 gene; *esr2b*: estrogen receptor 2b gene; *esr1*: estrogen receptor 1 gene; *cyp19a1b*: aromatase b gene.

### Seasonal differences in circulating steroids and in brain gene expression

A total of 8 circulating steroids were quantified in each season: A4, T, 11-KT, DHEA, E1, E2, progesterone and cortisol. Of this array of hormones, 5 were detectable in at least one season (Fig. 2). There were no detectable levels of 11-KT, DHEA or progesterone in any of the samples. A4 was significantly higher in the non-breeding season (Mann–Whitney U test, p = 0.0486), while T showed no seasonal difference (Mann–Whitney U test, p = 0.324). Cortisol was significantly higher in the breeding season (Mann–Whitney U test, p=0.0223). Both E1 and E2 exhibited seasonal patterns, being detectable only during the breeding season (E1: 4/15, 26.7%; E2: 5/15, 33.3%). To assess the significance of these seasonal differences in estrogen presence versus absence, we performed Fisher’s exact tests. In spite of being detectable only in the breeding season, E1 did not show a statistical significant variation between seasons (Fisher’s exact test, p = 0.113), whereas E2 displayed a statistical trend toward seasonal variation (Fisher’s exact test, p = 0.053).

**Figure 2:**
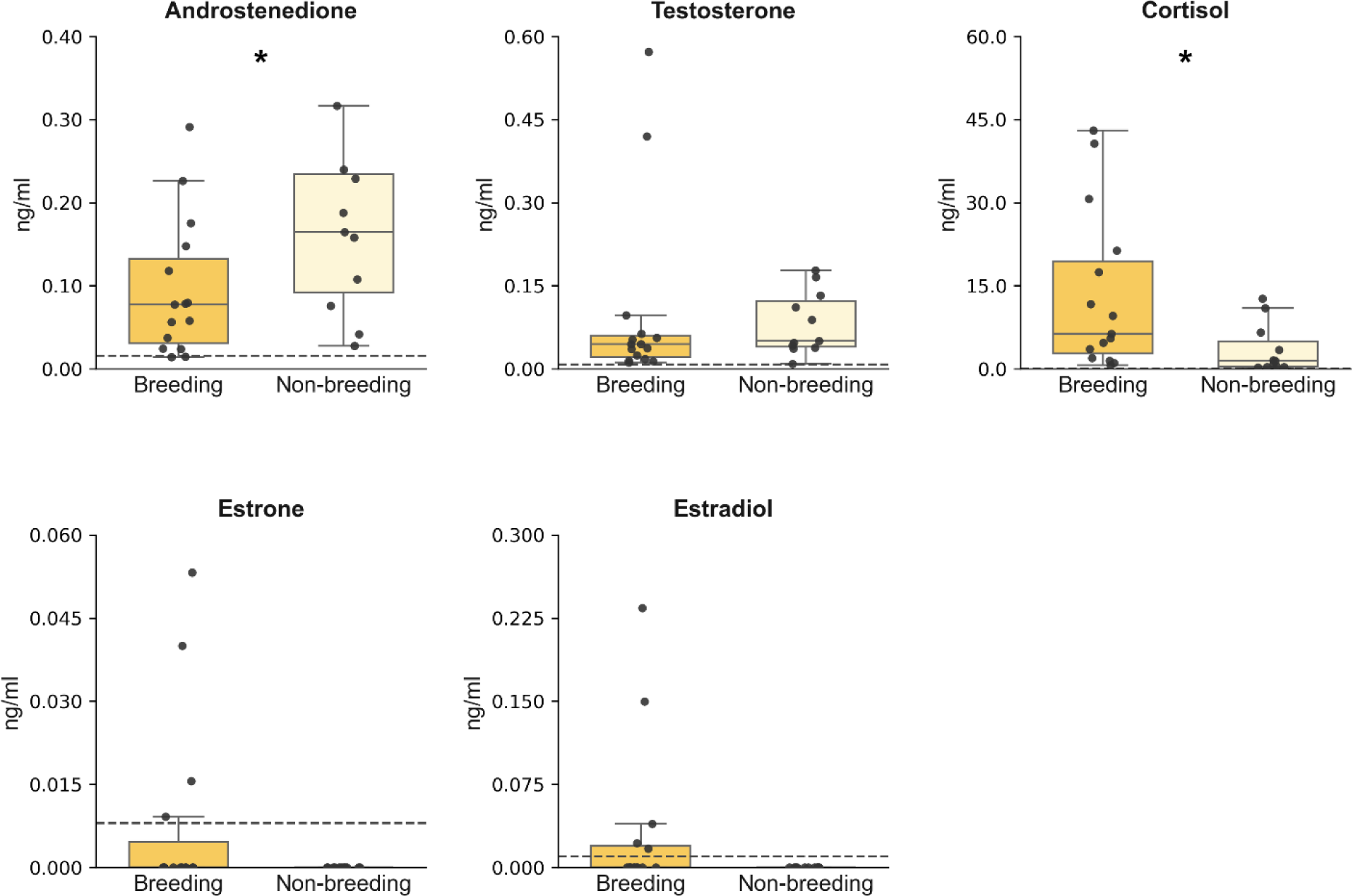
Female circulating steroid hormones in breeding and non-breeding seasons. A. Circulating androstenedione levels were higher during the non-breeding season, while cortisol levels were significantly higher in the breeding season. B. There was no significant difference in circulating estrone and estradiol levels between seasons. Note that estrone and estradiol were detectable only in some females during the breeding season. Dotted lines indicate the detection limit.

Females showed seasonal differences in GSI (breeding females: mean ± SD = 2.501 ± 2.414; non-breeding females: mean ± SD = 0.499 ± 0.236; p = 0.002). The unexpectedly low proportion of breeding females with detectable estrogen levels was independent of their gonadosomatic index (GSI), as no significant correlation was found (E1 ⍴ = 0.635, p_adjust = 0.663; E2: ⍴ = 0.203, p_adjust = 0.922).

In these same females, we quantified the expression of four genes (*cyp19a1b, esr1, esr2b* and *ar2*) in three regions of the SBN (POA, Vv and Vs). Gene expression varied seasonally in site-specific manner (Fig. 3). Non-breeding females showed a significantly higher *cyp19a1b* expression than breeding females in the Vv (Welch’s *t*-test, p = 0.0170) and the Vs (Welch’s *t*-test, p = 0.0242), whereas *cyp19a1b* expression was not different across seasons in the POA (Welch’s *t*-test, p = 0.937). One of the estrogen receptor genes, *esr2b*, was upregulated during the non-breeding season in the Vs (Welch’s *t*-test, p = 0.0011), but without seasonal differences in the POA (Welch’s *t*-test, p = 0.691) and in the Vv (Welch’s *t*-test, p = 0.110). Neither *esr1* nor *ar2* varied between seasons in any area (*esr1* POA: Welch’s *t*-test, p = 0.760; *esr1* Vv: Welch’s *t*-test, p = 0.136; *esr1* Vs: Welch’s *t*-test, p = 0.810; *ar2* POA: Mann–Whitney U test, p = 0.549; *ar2* Vv: Welch’s *t*-test, p = 0.585; *ar2* Vs: Welch’s *t*-test, p = 0.580).

**Figure 3:**
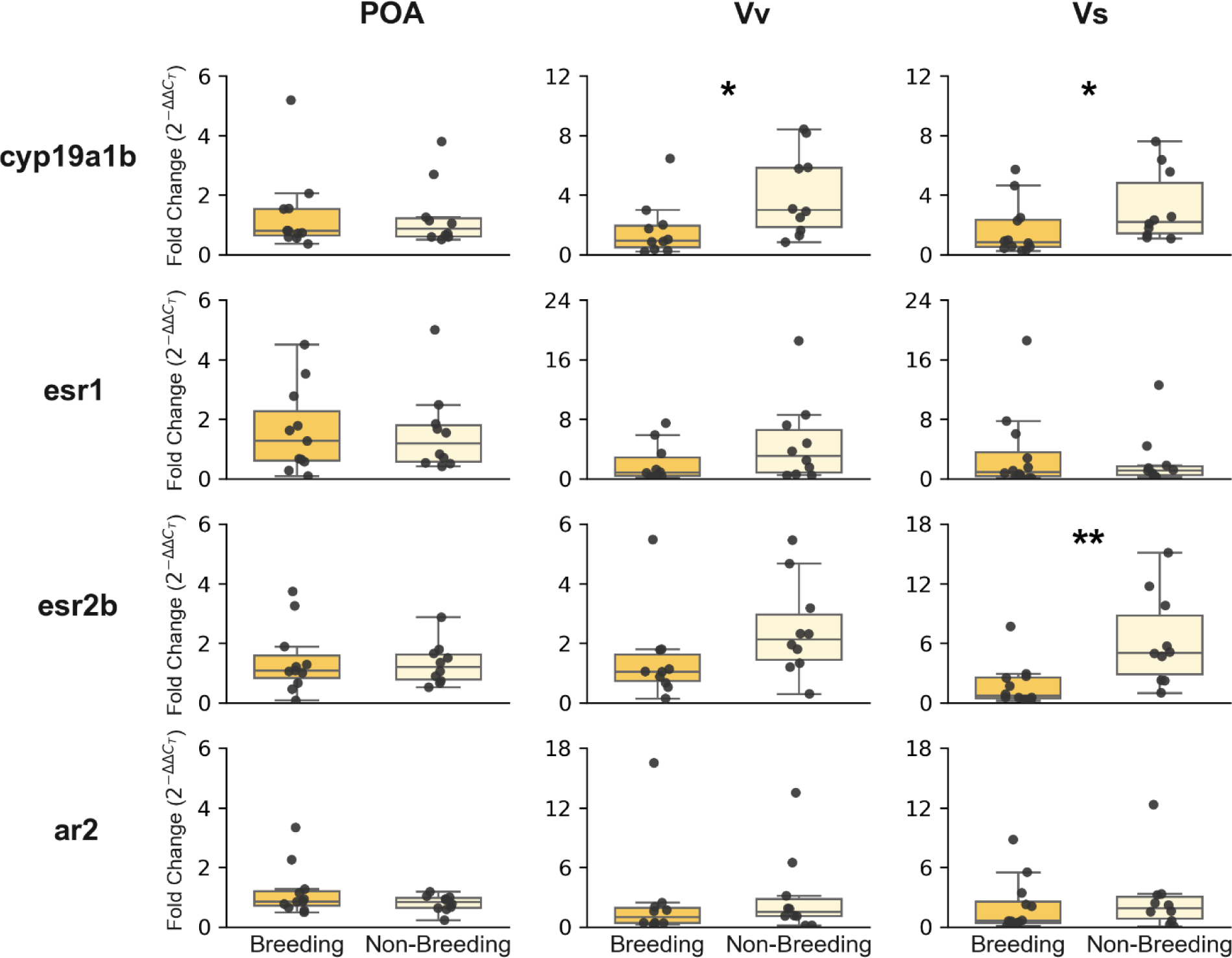
Quantification of gene expression in the social brain network of breeding and non-breeding females. *cyp19a1b* and *esr2b* expression were significantly higher in the Vv and Vs during the non-breeding season compared to the breeding season. Gene expression is shown as fold change relative to the breeding season (reference group). Abbreviations, POA: preoptic area; Vv: ventral nucleus of the ventral telencephalon; Vs: supracommissural nucleus of the ventral telencephalon; *cyp19a1b*: brain aromatase gene; *esr1*: estrogen receptor 1 gene; *esr2b*: estrogen receptor 2b gene; *ar2*: androgen receptor 2 gene.

In males we quantified the same steroids as in females. Of this array of hormones, 4 were detectable in at least one season (Fig. 4). There were no detectable levels of DHEA, E1, E2 or progesterone in any of the samples. A4 and T were higher during the non-breeding season (A4: Mann–Whitney U test, p = 0.0004; T: Mann–Whitney U test, p = 0.0036) while cortisol was higher during the breeding season (Mann–Whitney U test, p = 0.0362). 11-KT did not vary between seasons (Mann–Whitney U test, p = 0.0769). Males also showed an expected seasonal variation in their GSI (breeding males: mean ± SD = 0.285 ± 0.073; non-breeding males: mean ± SD = 0.102 ± 0.0789; p = 0.0002).

**Figure 4:**
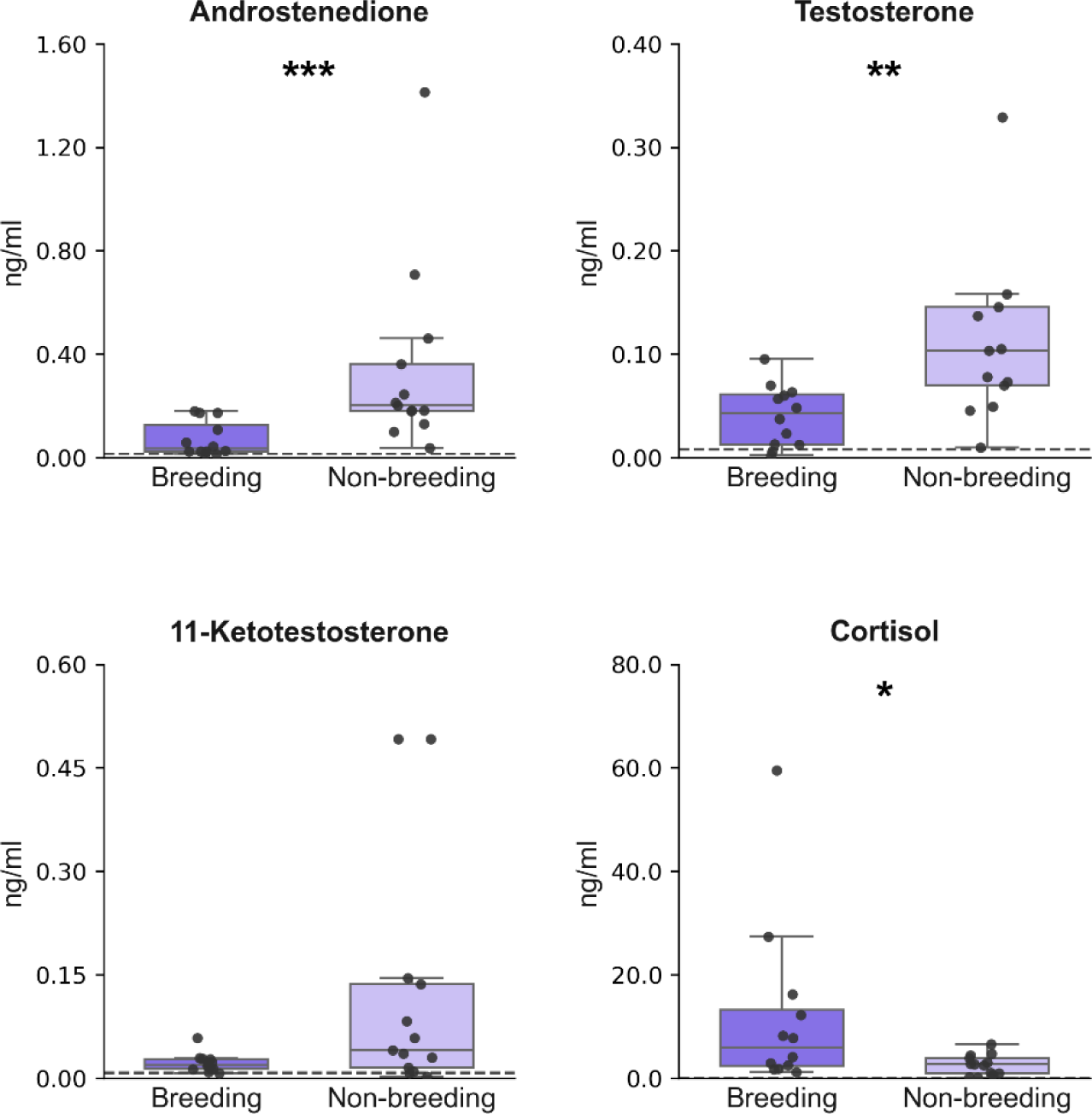
Male circulating steroid hormones in breeding and non-breeding season. Circulating androstenedione and testosterone levels were higher during the non-breeding season, while cortisol levels were significantly higher in the breeding season. Dotted lines indicate the detection limit.

In these same males, we quantified the expression of four genes in three brain areas of the SBN as mentioned for females. The expression of genes in the SBN varied between seasons and was site-specific (Fig. 5). *cyp19a1b* expression was higher in non-breeding males in the Vv (Welch’s *t*-test, p = 0.007) and in the Vs (Welch’s *t*-test, p = 0.00007), but not in the POA (Welch’s *t*-test, p = 0.895). Non-breeding males had more expression of both estrogen receptor genes in the Vs (esr1: Welch’s *t*-test, p = 0.006; esr2b: Welch’s *t*-test, p = 00002), but not in other areas (*esr1* POA: Welch’s *t*-test, p = 0.193; *esr1* Vv: Welch’s *t*-test, p = 0.128; *esr2b* POA: Welch’s *t*-test, p = 0.907; *esr2b* Vv: Welch’s *t*-test, p = 0.390). *ar2* did not vary between seasons in the POA (Mann– Whitney U test, p = 0.157), Vv (Welch’s *t*-test, p = 0.396) or Vs (Welch’s *t*-test, p = 0.069).

**Figure 5:**
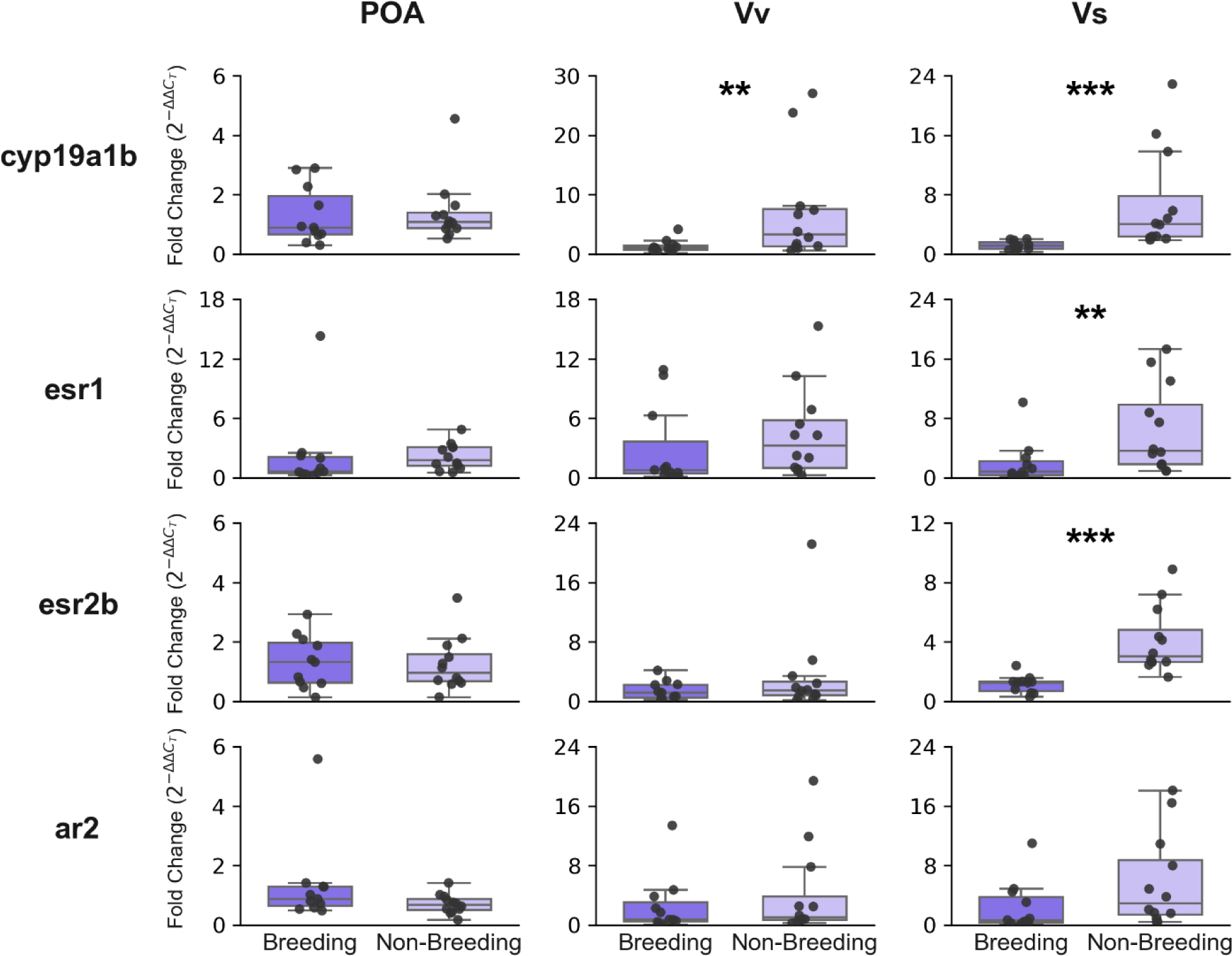
Quantification of gene expression in the social brain network of breeding and non-breeding males. *cyp19a1b* expression was significantly higher in the Vv and Vs during the non-breeding season. *esr1* and *esr2b expressions* were both significantly higher in the Vs during the non-breeding season. Gene expression is shown as fold change relative to the breeding season (reference group). POA: preoptic area; Vv: ventral nucleus of the ventral telencephalon; Vs: supracommissural nucleus of the ventral telencephalon; *cyp19a1b*: brain aromatase gene; *esr1*: estrogen receptor 1 gene; *esr2b*: estrogen receptor 2b gene; *ar2*: androgen receptor 2 gene.

### Sex differences in circulating steroids and in brain gene expression

Having established the seasonal patterns for each sex, we then examined whether females and males differed within each season. During the breeding season, there were sex differences in some of the quantified hormones; E1 and E2 were only present in breeding females, with E2 significantly higher in females (E1: Fischer exact test, p = 0.106; E2: Fischer exact test, p = 0.047), while 11-KT was only present in males (Fischer exact test, p = 0.0009). There were no significant differences in the other circulating hormones (A4: Mann–Whitney U test, p = 0.367; T: Mann– Whitney U test, p = 0.608; cortisol: Mann–Whitney U test, p = 0.788; Supp. Fig. 1). During the breeding season, at the transcriptional level, *esr1* was upregulated in the Vs of females (Welch’s *t*-test, p = 0.029) while no significant sex differences were observed for the other genes (POA *cyp19b1a*: Welch’s *t*-test, p = 0.776; POA *esr1*: Welch’s *t*-test, p = 0.686; POA *esr2b*: Welch’s *t*-test, p = 0.359; POA *ar2*: Mann–Whitney U, p = 0.694; Vv *cyp19b1a*: Welch’s *t*-test, p = 0.463; Vv *esr1*: Welch’s *t*-test, p = 0.765; Vv *esr2b*: Welch’s *t*-test, p = 0.404; Vv *ar2*: Welch’s *t*-test, p = 0.998; Vs *cyp19b1a*: Welch’s *t*-test, p = 0.175; Vs *esr2b*: Welch’s *t*-test, p = 0.862; Vs *ar2*: Welch’s *t*-test, p = 0.195; Supp. Fig. 2).

During the non-breeding season, sex differences were less pronounced (Supp. Fig. 3; Supp. Fig. 4). There was no significant difference in A4 (Mann–Whitney U test, p = 0.271), T (Mann– Whitney U test, p = 0.271), or cortisol (Mann–Whitney U test, p = 0.862). The only significant difference in circulating hormones was in 11-KT, which was present exclusively in males (Fischer exact test, p = 0.0002). At the transcriptional level, no significant sex differences were observed in any gene in the three analyzed nodes of the SBN (POA *cyp19b1a*: Welch’s *t*-test, p = 0.627; POA *esr1*: Welch’s *t*-test, p = 0.513; POA *esr2b*: Welch’s *t*-test, p = 0.490; POA *ar2*: Mann– Whitney U test, p = 0.621; Vv *cyp19b1a*: Welch’s *t*-test, p = 0.300; Vv *esr1*: Welch’s *t*-test, p = 0.760 Vv *esr2b*: Welch’s *t*-test, p = 0.805; Vv *ar2*: Welch’s *t*-test, p = 0.756; Vs *cyp19b1a*: Welch’s *t*-test, p = 0.688; Vs *esr1*: Welch’s *t*-test, p = 0.677; Vs *esr2b*: Welch’s *t*-test, p = 0.483; Vs *ar2*: Welch’s *t*-test, p = 0.972).

### Correlations between circulating hormones and SBN gene expression

To examine potential relationships between circulating hormones and gene expression in SBN nuclei, we performed Spearman correlations (Supp. Fig. 5). Across all four groups, circulating A4 showed strong positive correlations with circulating T (breeding females: ⍴ = 0.917, p_adjust = 0.039; breeding males: ⍴ = 0.893, p_adjust = 0.029; non-breeding females: ⍴ = 0.976, p_adjust = 0.0002; non-breeding males: ⍴ = 0.944, p_adjust = 0.0005). Additionally, in breeding females, both circulating T and A4 showed strong positive correlations with cyp19b1a gene expression in the Vv (T: ⍴ = 0.95, p_adjust = 0.013; A4: ⍴ = 0.917, p_adjust = 0.039). In this same group, E1 showed a statistical trend toward a positive correlation with cortisol (⍴ = 0.888, p_adjust = 0.053).

## DISCUSSION

The present study investigates seasonal modulations in hormonal pathways that may regulate year-round territorial aggression in wild male and female *G. omarorum*. We observed seasonal variation in circulating hormones and neuroendocrine pathways, supporting the idea that different mechanisms alternate in controlling the same behavior across seasons. During the non-breeding season, circulating levels of weak androgenic precursors were elevated. This increase coincided with a seasonal upregulation of the brain aromatase gene in the SBN, suggesting an enhanced capacity to convert these precursors into neuroestrogens. Simultaneously, we observed seasonally increased expression of estrogen receptor genes in the same network, potentially increasing neural sensitivity to these steroids and further amplifying the influence of neuroestrogens during the non-breeding season. In contrast, during the breeding season, circulating steroid levels were not elevated, indicating that breeding territorial aggression may rely on dynamic, stimulus-dependent mechanisms. Overall, this study provides the first evidence in teleost fish of seasonal plasticity in the mechanisms that can sustain year-round territorial aggression.

We examined the circulating levels of both A4 and DHEA, weak androgens with relatively low affinity for the androgen receptor [61,62]. A4 was detected in circulation in both sexes, with concentrations peaking during the non-breeding period. We propose that circulating A4 could serve as a precursor for local brain estrogen synthesis. This possibility is consistent with the high affinity of fish brain aromatase for A4 [23,63]. In addition, both males and females of *G. omarorum* show E1 in the forebrain (the direct aromatization product of A4) but not in circulation during the non-breeding season [51]. Reports on extragonadal sources of A4 in teleosts [64] and the fact that castrated *G. omarorum* maintain robust aggressive behavior in the non-breeding season [16] suggest that during this period A4 may derive partly from non-gonadal tissues. However, direct evidence for the role of A4 in modulating aggression is lacking and would require measurements of brain A4 uptake and estrogen conversion rates. Interestingly, in mammals and some birds displaying non-breeding aggression, several studies have reported higher DHEA concentrations during the non-breeding season compared to the breeding season [30–34], raising the possibility that this steroid contributes to year-round territorial aggression, though notable exceptions to the DHEA pattern exist in certain avian species [59,65,66]. In *G. omarorum*, DHEA was undetectable across seasons and sexes. This pattern suggests that in this species either androgen synthesis may bypass DHEA, or alternatively, that DHEA is rapidly metabolized to other steroids. From an evolutionary perspective, the recruitment of weak circulating androgens as a neurosteroid precursor in *G. omarorum* could indicate a broad pattern across vertebrates, where weak androgen precursors sustain aggression beyond the breeding season through local aromatization or conversion into active androgens. In *G. omarorum*, however, the weak precursor implicated differs from those in mammals and birds, suggesting lineage-specific solutions to sustaining non-breeding aggression.

The role of estrogens in aggression is well documented in fish [67]. Our results on the seasonal analysis of the expression of *cyp19a1b* revealed consistent results across sexes and brain sites. In both males and females, there is an upregulation of this gene during the non-breeding season in the ventral nucleus of the ventral telencephalon (Vv) and the ventral supracommissural nucleus (Vs), but not in the preoptic area (POA). Overall, these results align with a seasonal increase in the brain’s capacity to synthesize estrogens, most probably from circulating precursors, in specific aggression-related areas. Estrogens are known modulators of non-breeding aggressive behavior in *G. omarorum* [16,19]. However, while no circulating estrogens are detectable during this season, estrogens are detectable in the forebrains of both males and females [51]. The results of the present study suggest that at least part of this brain estrogen is produced within the SBN to sustain non-breeding aggression in both males and females. Patterns of aromatase expression vary across vertebrates: unlike *G. omarorum*, song sparrows show no aromatase overexpression during the non-breeding season [25,68], whereas in Siberian hamsters, expression is site and sex-dependent, with some areas upregulating and others downregulating aromatase during the non-breeding season [40]. These results underscore the importance of additional research in different species to identify evolutionary patterns.

The potential role of neuroestrogens in modulating non-breeding territorial aggression in *G. omarorum* is also supported by the seasonal upregulation of *esr1* and *esr2b* in nuclei of the SBN, which likely increases the sensitivity of neural circuits to estrogenic signaling. In lab settings, acute pharmacological inhibition of estrogen synthesis rapidly reduces both the motivation to engage and the intensity of aggressive behavior in male–male and female–female dyads during the non-breeding season [16,19], indicating rapid, non-genomic effects mediated by estrogen receptors [69,70]. Furthermore, these receptors can also act more slowly through genomic mechanisms by regulating target gene transcription [71]. For instance, in teleost fish, *cyp19a1b* transcription is driven by estrogen response elements [72], which might indicate that upregulated *esr1* and *esr2b* in the SBN of *G. omarorum* could be maintaining the elevated aromatase expression shown during the non-breeding season. These modes of action are not mutually exclusive and can interact [73]. The specific contributions of each estrogen receptor subtype during this period in *G. omarorum* remain to be clarified. Similar seasonal modulation of brain estrogen sensitivity occurs in other species with year-round territorial aggression, including mammals [18,38] and birds [36], though this pattern is not universal across all taxa [68,74,75], suggesting that even among species with year-round territorial aggression, different neuroendocrine mechanisms can sustain similar behavioral phenotypes.

In addition to neuroestrogens, androgens also modulate non-breeding territorial aggression in *G. omarorum* through direct activation of androgen receptors in both sexes [20]. We show that potent androgens are detectable in circulation during the non-breeding season in both sexes, as shown in previous reports in *G. omarorum* [16,51]. This is consistent with our finding of *ar2* expression in the SBN during the non-breeding season. Both 11-KT and T may bind to androgen receptors, although the latter could also serve as a substrate for local estrogen synthesis, as discussed above. We found no seasonal differences in *ar2* expression in males or females, suggesting that androgen receptors may also contribute to territorial aggression during the breeding season.

Our results on breeding circulating steroid hormones indicate that territorial and reproductive behaviors, including territorial aggression and reproduction, can persist even under low hormonal baselines. We propose that in *G. omarorum*, sex steroids rise rapidly and transiently in response to social challenges or reproductive opportunities, rather than remaining consistently high across the breeding season. This dynamic pattern aligns with the challenge hypothesis [76,77], which has been reported in most teleost species examined, in contrast to other vertebrate taxa where evidence is less consistent (reviewed in [78]). Such flexibility would allow individuals to increase hormone levels associated with competitive behaviors only when needed, while minimizing the physiological costs of prolonged steroid elevation, such as immunosuppression [79] and increased metabolic expenditure [80]. The absence of sexual dimorphism in *G. omarorum*, both morphologically and in electric signal characteristics [81,82], further supports the idea that androgen action during breeding is transient rather than constitutive, as there are no secondary sexual traits that need to be maintained by increased basal hormone levels. In line with this, in males of the weakly electric gymnotiform *Brachyhypopomus gauderio*, which displays seasonal sexual dimorphism both in its morphology and different aspects of its electric signaling, there is a pronounced seasonal variation in circulating levels of 11-KT [83] and these secondary sexual traits are inducible by androgen treatments [84,85]. In females of *G. omarorum*, estrogens were detected exclusively during the breeding season in a fraction of the population, likely reflecting estrogen-dependent vitellogenesis [86]. However, we cannot rule out a role of neuroestrogens in female breeding aggression, supported by the positive correlations between A4 and T and brain aromatase expression in the Vv. Overall, our findings suggest that *G. omarorum* relies on flexible, short-term endocrine responses to balance the demands of reproduction, parental investment, and territorial defense, rather than on chronically elevated sexual hormones. Nonetheless, the potential role of sex steroids and their dynamics in modulating breeding aggression remains to be tested through pharmacological inhibition and post-interaction steroid measurements.

Cortisol plays a central role in the stress physiology of fish, mediating the redistribution of energetic resources needed to cope with social and environmental challenges [87]. Most research on stress in fish has been conducted in captive or cultured populations, where animals are exposed to artificial or chronic stressors, limiting the understanding of how stress functions under natural conditions [88]. By examining free-living animals, our study shows that in both sexes basal cortisol levels are higher during the breeding season in *G. omarorum*. This elevation may reflect the greater energetic demands of reproduction, consistent with findings in vertebrates [89], including fish species, where cortisol levels often peak during the breeding season [90,91]. Beyond energetic mobilization, elevated cortisol during breeding may also regulate social behaviors; in the electric fish *Apteronotus leptorhynchus*, cortisol implants increased aggressive electrocommunication signals [92], suggesting cortisol can directly modulate breeding-season social interactions.

Our results show that neural circuits undergo distinct seasonal transitions in each sex. In the Vs, males overexpress both esr1 and esr2b during the non-breeding season, while females overexpress only esr2b. These sexually dimorphic trajectories may be viewed as compensatory mechanisms (inspired by the compensation theory, [93]), counterbalancing physiological disparities and converging toward a common SBN configuration in the non-breeding season when the motivation to fight over territory is expected to be similar across sexes [46], aggressive displays are sexually monomorphic [42,48], and neuroestrogens modulate both male and female aggression [16,19]. These results position *G. omarorum* as a valuable model for studying seasonal transitions in neural configuration.

### Conclusions

We found evidence of seasonal plasticity in the hormonal and neuroendocrine mechanisms underlying sustained year-round territorial aggression in females and males of *G. omarorum*. Specifically, we documented seasonal shifts in circulating hormone levels coupled with changes in brain aromatase and estrogen receptor expression within the SBN, suggesting that distinct hormonal configurations may alternately control persistent aggressive behavior across seasons. Species that maintain year-round territorial aggression provide excellent models to understand how consistent behavioral outputs can arise from flexible neural mechanisms, offering a framework to explore how animals adapt their physiology to environmental and social demands. Originally proposed in birds and mammals [27–29], the idea of seasonally alternating mechanisms underlying year-round territorial aggression is extended here to teleost fish. This suggests that neuroendocrine plasticity may represent a widespread mechanism for maintaining stable behavioral phenotypes across variable, but predictable environments.

## BIBLIOGRAPHY

1. Tramontin AD, Brenowitz EA. 2000 Seasonal plasticity in the adult brain. Trends in Neurosciences 23, 251–258. (doi:10.1016/S0166-2236(00)01558-7)

2. Piersma T, Drent J. 2003 Phenotypic flexibility and the evolution of organismal design. Trends in Ecology & Evolution 18, 228–233. (doi:10.1016/S0169-5347(03)00036-3)

3. Newman SW. 1999 The Medial Extended Amygdala in Male Reproductive Behavior A Node in the Mammalian Social Behavior Network. Annals of the New York Academy of Sciences 877, 242–257. (doi:10.1111/j.1749-6632.1999.tb09271.x)

4. Goodson JL, Evans AK, Soma KK. 2005 Neural responses to aggressive challenge correlate with behavior in nonbreeding sparrows. NeuroReport 16, 1719–1723. (doi:10.1097/01.wnr.0000183898.47160.15)

5. Kelly AM, Vitousek MN. 2017 Dynamic modulation of sociality and aggression: an examination of plasticity within endocrine and neuroendocrine systems. Phil. Trans. R. Soc. B 372, 20160243. (doi:10.1098/rstb.2016.0243)

6. Nelson RJ. 2005 Biology of Aggression. Oxford University Press.

7. Wingfield JC, Moore IT, Goymann W, Wacker DW, Sperry TS. 2006 Contexts and ethology of vertebrate aggression: implications for the evolution of hormone-behavior interactions. In Biology of aggression, pp. 179–210.

8. Soma KK. 2006 Testosterone and Aggression: Berthold, Birds and Beyond. J Neuroendocrinology 18, 543–551. (doi:10.1111/j.1365-2826.2006.01440.x)

9. Nelson RJ, Trainor BC. 2007 Neural mechanisms of aggression. Nat Rev Neurosci 8, 536–546. (doi:10.1038/nrn2174)

10. Wu MV, Manoli DS, Fraser EJ, Coats JK, Tollkuhn J, Honda S-I, Harada N, Shah NM. 2009 Estrogen Masculinizes Neural Pathways and Sex-Specific Behaviors. Cell 139, 61–72. (doi:10.1016/j.cell.2009.07.036)

11. Rubenstein DR, Wikelski M. 2005 Steroid hormones and aggression in female Galápagos marine iguanas. Hormones and Behavior 48, 329–341. (doi:10.1016/j.yhbeh.2005.04.006)

12. Duque-Wilckens N, Trainor BC, Marler CA. 2019 Aggression and Territoriality. In Encyclopedia of Animal Behavior, pp. 539–546. Elsevier. (doi:10.1016/B978-0-12-809633-8.90064-5)

13. Rosvall KA, Bentz AB, George EM. 2020 How research on female vertebrates contributes to an expanded challenge hypothesis. Hormones and Behavior 123, 104565. (doi:10.1016/j.yhbeh.2019.104565)

14. Wingfield, John C. 1994 Regulation of territorial behavior in the sedentary song sparrow, Melospiza melodia morphna. Hormones and Behavior 28.

15. Demas GE, Moffatt CA, Drazen DL, Nelson RJ. 1999 Castration Does Not Inhibit Aggressive Behavior in Adult Male Prairie Voles (Microtus ochrogaster). Physiology & Behavior 66, 59–62. (doi:10.1016/S0031-9384(98)00268-6)

16. Jalabert C, Quintana L, Pessina P, Silva A. 2015 Extra-gonadal steroids modulate non-breeding territorial aggression in weakly electric fish. Hormones and Behavior 72, 60–67. (doi:10.1016/j.yhbeh.2015.05.003)

17. Soma KK, Sullivan K, Wingfield J. 1999 Combined Aromatase Inhibitor and Antiandrogen Treatment Decreases Territorial Aggression in a Wild Songbird during the Nonbreeding Season. General and Comparative Endocrinology 115, 442–453. (doi:10.1006/gcen.1999.7334)

18. Trainor BC, Lin S, Finy* MS, Rowland MR, Nelson RJ. 2007 Photoperiod reverses the effects of estrogens on male aggression via genomic and nongenomic pathways. Proc. Natl. Acad. Sci. U.S.A. 104, 9840–9845. (doi:10.1073/pnas.0701819104)

19. Zubizarreta L, Silva AC, Quintana L. 2020 The estrogenic pathway modulates non-breeding female aggression in a teleost fish. Physiology & Behavior 220, 112883. (doi:10.1016/j.physbeh.2020.112883)

20. Valiño G, Dunlap K, Quintana L. 2024 Androgen receptors rapidly modulate non-breeding aggression in male and female weakly electric fish (Gymnotus omarorum). Hormones and Behavior 159, 105475. (doi:10.1016/j.yhbeh.2023.105475)

21. Compagnone NA, Mellon SH. 2000 Neurosteroids: Biosynthesis and Function of These Novel Neuromodulators. Frontiers in Neuroendocrinology 21, 1–56. (doi:10.1006/frne.1999.0188)

22. Tsutsui K, Ukena K, Usui M, Sakamoto H, Takase M. 2000 Novel brain function: biosynthesis and actions of neurosteroids in neurons. Neuroscience Research 36, 261–273. (doi:10.1016/S0168-0102(99)00132-7)

23. Diotel N, Do Rego J-L, Anglade I, Vaillant C, Pellegrini E, Vaudry H, Kah O. 2011 The Brain of Teleost Fish, a Source, and a Target of Sexual Steroids. Front. Neurosci. 5. (doi:10.3389/fnins.2011.00137)

24. Do Rego JL, Vaudry H. 2016 Comparative aspects of neurosteroidogenesis: From fish to mammals. General and Comparative Endocrinology 227, 120–129. (doi:10.1016/j.ygcen.2015.05.014)

25. Soma KK, Schlinger BA, Wingfield JC, Saldanha CJ. 2003 Brain aromatase, 5α-reductase, and 5β-reductase change seasonally in wild male song sparrows: Relationship to aggressive and sexual behavior. J. Neurobiol. 56, 209–221. (doi:10.1002/neu.10225)

26. Pradhan DS, Newman AEM, Wacker DW, Wingfield JC, Schlinger BA, Soma KK. 2010 Aggressive interactions rapidly increase androgen synthesis in the brain during the non-breeding season. Hormones and Behavior 57, 381–389. (doi:10.1016/j.yhbeh.2010.01.008)

27. Wingfield JC, Lynn SE, Soma KK. 2001 Avoiding the ‘Costs’ of Testosterone: Ecological Bases of Hormone-Behavior Interactions. Brain Behav Evol 57, 239–251. (doi:10.1159/000047243)

28. Soma KK, Rendon NM, Boonstra R, Albers HE, Demas GE. 2015 DHEA effects on brain and behavior: Insights from comparative studies of aggression. The Journal of Steroid Biochemistry and Molecular Biology 145, 261–272. (doi:10.1016/j.jsbmb.2014.05.011)

29. Demas GE, Munley KM, Jasnow AM. 2023 A seasonal switch hypothesis for the neuroendocrine control of aggression. Trends in Endocrinology & Metabolism 34, 799–812. (doi:10.1016/j.tem.2023.08.015)

30. Hau M, Stoddard ST, Soma KK. 2004 Territorial aggression and hormones during the non-breeding season in a tropical bird. Hormones and Behavior 45, 40–49. (doi:10.1016/j.yhbeh.2003.08.002)

31. González-Gómez PL, Blakeslee WS, Razeto-Barry P, Borthwell RM, Hiebert SM, Wingfield JC. 2014 Aggression, body condition, and seasonal changes in sex-steroids in four hummingbird species. J Ornithol 155, 1017–1025. (doi:10.1007/s10336-014-1088-y)

32. Rendon NM, Rudolph LM, Sengelaub DR, Demas GE. 2015 The agonistic adrenal: melatonin elicits female aggression via regulation of adrenal androgens. Proc. R. Soc. B. 282, 20152080. (doi:10.1098/rspb.2015.2080)

33. Boonstra R, Dušek A, Flynn RW. 2018 DHEA and territoriality during the nonbreeding season in male American martens (Martes americana). Journal of Mammalogy 99, 826–835. (doi:10.1093/jmammal/gyy067)

34. Boonstra R, Gandhi N, Kraushaar A, Galbreath K. 2022 From Habitat to Hormones: Year-around territorial behavior in rock-dwelling but not in forest and grassland lagomorphs and the role of DHEA. Hormones and Behavior 142, 105179. (doi:10.1016/j.yhbeh.2022.105179)

35. Jalabert C, Gray SL, Soma KK. 2024 An Aggressive Interaction Rapidly Increases Brain Androgens in a Male Songbird during the Non-breeding Season. J. Neurosci. 44, e1095232024. (doi:10.1523/JNEUROSCI.1095-23.2024)

36. Canoine V, Fusani L, Schlinger B, Hau M. 2007 Low sex steroids, high steroid receptors: Increasing the sensitivity of the nonreproductive brain. Developmental Neurobiology 67, 57–67. (doi:10.1002/dneu.20296)

37. Trainor BC, Rowland MR, Nelson RJ. 2007 Photoperiod affects estrogen receptor α, estrogen receptor β and aggressive behavior. Eur J of Neuroscience 26, 207–218. (doi:10.1111/j.1460-9568.2007.05654.x)

38. Kramer KM, Simmons JL, Freeman DA. 2008 Photoperiod alters central distribution of estrogen receptor α in brain regions that regulate aggression. Hormones and Behavior 53, 358–365. (doi:10.1016/j.yhbeh.2007.11.002)

39. Rendon NM, Amez AC, Proffitt MR, Bauserman ER, Demas GE. 2017 Aggressive behaviours track transitions in seasonal phenotypes of female Siberian hamsters. Functional Ecology 31, 1071–1081. (doi:10.1111/1365-2435.12816)

40. Munley KM, Sinkiewicz DM, Szwed SM, Demas GE. 2023 Sex and seasonal differences in neural steroid sensitivity predict territorial aggression in Siberian hamsters. Hormones and Behavior 154, 105390. (doi:10.1016/j.yhbeh.2023.105390)

41. Silva AC, Perrone R, Zubizarreta L, Batista G, Stoddard PK. 2013 Neuromodulation of the agonistic behavior in two species of weakly electric fish that display different types of aggression. Journal of Experimental Biology 216, 2412–2420. (doi:10.1242/jeb.082180)

42. Quintana L, Zubizarreta L, Jalabert C, Batista G, Perrone R, Silva A. 2016 Building the case for a novel teleost model of non-breeding aggression and its neuroendocrine control. Journal of Physiology-Paris 110, 224–232. (doi:10.1016/j.jphysparis.2016.11.009)

43. Silva AC, Zubizarreta L, Quintana L. 2020 A Teleost Fish Model to Understand Hormonal Mechanisms of Non-breeding Territorial Behavior. Front. Endocrinol. 11, 468. (doi:10.3389/fendo.2020.00468)

44. Quintana L, Jalabert C, Fokidis HB, Soma KK, Zubizarreta L. 2021 Neuroendocrine Mechanisms Underlying Non-breeding Aggression: Common Strategies Between Birds and Fish. Front. Neural Circuits 15, 716605. (doi:10.3389/fncir.2021.716605)

45. Quintana L, Salazar V. 2025 Social behavior in South American electric fishes: Linking neuroendocrine regulation, signal plasticity, and reproductive strategies. Neuroscience 573, 154–166. (doi:10.1016/j.neuroscience.2025.03.028)

46. Zubizarreta L, Quintana L, Hernández D, Teixeira De Mello F, Meerhoff M, Massaaki Honji R, Guimarães Moreira R, Silva A. 2020 Seasonal and social factors associated with spacing in a wild territorial electric fish. PLoS ONE 15, e0228976. (doi:10.1371/journal.pone.0228976)

47. Migliaro A, Pedraja F, Mucha S, Benda J, Silva A. 2025 Tracking spatial patterns and daily modulation of behavior in a natural population of the pulse-type weakly electric fish, Gymnotus omarorum. iScience 28, 112018. (doi:10.1016/j.isci.2025.112018)

48. Batista G, Zubizarreta L, Perrone R, Silva A. 2012 Non-sex-biased Dominance in a Sexually Monomorphic Electric Fish: Fight Structure and Submissive Electric Signalling. Ethology 118, 398–410. (doi:10.1111/j.1439-0310.2012.02022.x)

49. Zubizarreta L, Stoddard PK, Silva A. 2015 Aggression Levels Affect Social Interaction in the Non-Breeding Territorial Aggression of the Weakly Electric Fish, *Gymnotus omarorum*. Ethology 121, 8–16. (doi:10.1111/eth.12299)

50. Perrone R, Pedraja F, Valiño G, Tassino B, Silva A. 2019 Non-breeding territoriality and the effect of territory size on aggression in the weakly electric fish, Gymnotus omarorum. acta ethol 22, 79–89. (doi:10.1007/s10211-019-00309-7)

51. Zubizarreta L, Jalabert C, Silva AC, Soma KK, Quintana L. 2023 Brain and circulating steroids in an electric fish: Relevance for non-breeding aggression. PLoS ONE 18, e0289461. (doi:10.1371/journal.pone.0289461)

52. Migliaro A, Moreno V, Marchal P, Silva A. 2018 Daily changes in the electric behavior of weakly electric fish in nature persist in constant darkness and are socially synchronized. Biology Open, bio.036319. (doi:10.1242/bio.036319)

53. Silva A, Quintana L, Galeano M, Errandonea P. 2003 Biogeography and Breeding in Gymnotiformes from Uruguay. Environmental Biology of Fishes 66, 329–338. (doi:10.1023/A:1023986600069)

54. Pouso P, Radmilovich M, Silva A. 2017 An immunohistochemical study on the distribution of vasotocin neurons in the brain of two weakly electric fish, Gymnotus omarorum and Brachyhypopomus gauderio. Tissue and Cell 49, 257–269. (doi:10.1016/j.tice.2017.02.003)

55. Palkovits M. 1979 Microchemistry of Microdissected Hypothalamic Nuclear Areas. In International Review of Cytology, pp. 315–339. Elsevier. (doi:10.1016/S0074-7696(08)61825-2)

56. Eastman G, Valiño G, Radío S, Young RL, Quintana L, Zakon HH, Hofmann HA, Sotelo-Silveira J, Silva A. 2020 Brain transcriptomics of agonistic behaviour in the weakly electric fish Gymnotus omarorum, a wild teleost model of non-breeding aggression. Sci Rep 10, 9496. (doi:10.1038/s41598-020-66494-9)

57. Livak KJ, Schmittgen TD. 2001 Analysis of Relative Gene Expression Data Using Real-Time Quantitative PCR and the 2−ΔΔCT Method. Methods 25, 402–408. (doi:10.1006/meth.2001.1262)

58. Wei R, Wang J, Jia E, Chen T, Ni Y, Jia W. 2018 GSimp: A Gibbs sampler based left-censored missing value imputation approach for metabolomics studies. PLoS Comput Biol 14, e1005973. (doi:10.1371/journal.pcbi.1005973)

59. Jalabert C, Ma C, Soma KK. 2021 Profiling of systemic and brain steroids in male songbirds: Seasonal changes in neurosteroids. J Neuroendocrinology 33, e12922. (doi:10.1111/jne.12922)

60. Anderson MJ. 2001 A new method for non-parametric multivariate analysis of variance. Austral Ecology 26, 32–46. (doi:10.1111/j.1442-9993.2001.01070.pp.x)

61. Chen F et al. 2005 Direct Agonist/Antagonist Functions of Dehydroepiandrosterone. Endocrinology 146, 4568–4576. (doi:10.1210/en.2005-0368)

62. Jørgensen A, Andersen O, Bjerregaard P, Rasmussen LJ. 2007 Identification and characterisation of an androgen receptor from zebrafish Danio rerio. Comparative Biochemistry and Physiology Part C: Toxicology & Pharmacology 146, 561–568. (doi:10.1016/j.cbpc.2007.07.002)

63. González A, Piferrer F. 2002 Characterization of aromatase activity in the sea bass: effects of temperature and different catalytic properties of brain and ovarian homogenates and microsomes. J. Exp. Zool. 293, 500–510. (doi:10.1002/jez.90005)

64. Mayer I, Borg B, Schulz R. 1990 Seasonal changes in and effect of castration/androgen replacement on the plasma levels of five androgens in the male three-spined stickleback, Gasterosteus aculeatus L. General and Comparative Endocrinology 79, 23–30. (doi:10.1016/0016-6480(90)90084-Y)

65. Pintér O, Péczely P, Zsebõk S, Zelena D. 2011 Seasonal changes in courtship behavior, plasma androgen levels and in hypothalamic aromatase immunoreactivity in male free-living European starlings (Sturnus vulgaris). General and Comparative Endocrinology 172, 151–157. (doi:10.1016/j.ygcen.2011.02.002)

66. Landys MM, Goymann W, Soma KK, Slagsvold T. 2013 Year-round territorial aggression is independent of plasma DHEA in the European nuthatch Sitta europaea. Hormones and Behavior 63, 166–172. (doi:10.1016/j.yhbeh.2012.10.002)

67. Stennette KA, Godwin JR. 2024 Estrogenic influences on agonistic behavior in teleost fishes. Hormones and Behavior 161, 105519. (doi:10.1016/j.yhbeh.2024.105519)

68. Wacker DW, Wingfield JC, Davis JE, Meddle SL. 2010 Seasonal changes in aromatase and androgen receptor, but not estrogen receptor mRNA expression in the brain of the free-living male song sparrow, Melospiza melodia morphna. J of Comparative Neurology 518, 3819–3835. (doi:10.1002/cne.22426)

69. Boonyaratanakornkit V, Edwards D. 2007 Receptor Mechanisms Mediating Non-Genomic Actions of Sex Steroids. Semin Reprod Med 25, 139–153. (doi:10.1055/s-2007-973427)

70. Laredo SA, Villalon Landeros R, Trainor BC. 2014 Rapid effects of estrogens on behavior: Environmental modulation and molecular mechanisms. Frontiers in Neuroendocrinology 35, 447–458. (doi:10.1016/j.yfrne.2014.03.005)

71. Nelson ER, Habibi HR. 2013 Estrogen receptor function and regulation in fish and other vertebrates. General and Comparative Endocrinology 192, 15–24. (doi:10.1016/j.ygcen.2013.03.032)

72. Tchoudakova A, Kishida M, Wood E, Callard GV. 2001 Promoter characteristics of two cyp19 genes differentially expressed in the brain and ovary of teleost fish. The Journal of Steroid Biochemistry and Molecular Biology 78, 427–439. (doi:10.1016/S0960-0760(01)00120-0)

73. Wilkenfeld SR, Lin C, Frigo DE. 2018 Communication between genomic and non-genomic signaling events coordinate steroid hormone actions. Steroids 133, 2–7. (doi:10.1016/j.steroids.2017.11.005)

74. Trainor BC, Sima Finy M, Nelson RJ. 2008 Rapid effects of estradiol on male aggression depend on photoperiod in reproductively non-responsive mice. Hormones and Behavior 53, 192–199. (doi:10.1016/j.yhbeh.2007.09.016)

75. Villavicencio CP, Windley H, D’Amelio PB, Gahr M, Goymann W, Quispe R. 2021 Neuroendocrine patterns underlying seasonal song and year-round territoriality in male black redstarts. Front Zool 18, 8. (doi:10.1186/s12983-021-00389-x)

76. Wingfield JC, Hegner RE, Dufty, AM, Ball GF. 1990 The ‘Challenge Hypothesis’: Theoretical Implications for Patterns of Testosterone Secretion, Mating Systems, and Breeding Strategies. The American Naturalist 136, 829–846. (doi:10.1086/285134)

77. Goymann W, Moore IT, Oliveira RF. 2019 Challenge Hypothesis 2.0: A Fresh Look at an Established Idea. BioScience 69, 432–442. (doi:10.1093/biosci/biz041)

78. Moore IT, Hernandez J, Goymann W. 2020 Who rises to the challenge? Testing the Challenge Hypothesis in fish, amphibians, reptiles, and mammals. Hormones and Behavior 123, 104537. (doi:10.1016/j.yhbeh.2019.06.001)

79. Gubbels Bupp MR, Jorgensen TN. 2018 Androgen-Induced Immunosuppression. Front. Immunol. 9, 794. (doi:10.3389/fimmu.2018.00794)

80. Ros AFH, Becker K, Canário AVM, Oliveira RF. 2004 Androgen levels and energy metabolism in *Oreochromis mossambicus*. Journal of Fish Biology 65, 895–905. (doi:10.1111/j.0022-1112.2004.00484.x)

81. Richer-de-Forges MM, Crampton WGR, Albert JS. 2009 http://www.jstor.org A New Species of Gymnotus (Gymnotiformes, Gymnotidae) from Uruguay: Description of a Model Species in Neurophysiological Research. Copeia 2009, 538–544.

82. Craig JM, Kim LY, Tagliacollo VA, Albert JS. 2019 Phylogenetic revision of Gymnotidae (Teleostei: Gymnotiformes), with descriptions of six subgenera. PLoS ONE 14, e0224599. (doi:10.1371/journal.pone.0224599)

83. Dunlap KD, Silva AC, Chung M. 2011 Environmental complexity, seasonality and brain cell proliferation in a weakly electric fish, *Brachyhypopomus gauderio*. Journal of Experimental Biology 214, 794–805. (doi:10.1242/jeb.051037)

84. Silva A, Quintana L, Ardanaz JL, Macadar O. 2002 Environmental and hormonal influences upon EOD waveform in gymnotiform pulse fish. Journal of Physiology-Paris 96, 473–484. (doi:10.1016/S0928-4257(03)00003-2)

85. Goldina A, Gavassa S, Stoddard PK. 2011 Testosterone and 11-ketotestosterone have different regulatory effects on electric communication signals of male Brachyhypopomus gauderio. Hormones and Behavior 60, 139–147. (doi:10.1016/j.yhbeh.2011.03.014)

86. Reading BJ, Sullivan CV. 2024 Vitellogenesis in fishes. In Encyclopedia of Fish Physiology, pp. 626–636.

87. Schreck CB, Tort L. 2016 The concept of stress in fish. In Fish physiology, pp. 1–34.

88. Pankhurst NW. 2016 Reproduction and development. In Fish physiology.

89. Michael Romero L. 2002 Seasonal changes in plasma glucocorticoid concentrations in free-living vertebrates. General and Comparative Endocrinology 128, 1–24. (doi:10.1016/S0016-6480(02)00064-3)

90. Berg OK, Finstad. 2008 Energetic trade-off in reproduction: cost benefit considerations and plasticity. In Fish reproduction.

91. Cowan M, Azpeleta C, López-Olmeda JF. 2017 Rhythms in the endocrine system of fish: a review. J Comp Physiol B 187, 1057–1089. (doi:10.1007/s00360-017-1094-5)

92. Dunlap KD, Pelczar PL, Knapp R. 2002 Social Interactions and Cortisol Treatment Increase the Production of Aggressive Electrocommunication Signals in Male Electric Fish, Apteronotus leptorhynchus. Hormones and Behavior 42, 97–108. (doi:10.1006/hbeh.2002.1807)

93. De Vries GJ. 2004 Minireview: Sex Differences in Adult and Developing Brains: Compensation, Compensation, Compensation. Endocrinology 145, 1063–1068. (doi:10.1210/en.2003-1504)

