## Supplemental Figures for "Seasonal plasticity in neuroendocrine mechanisms relevant to year-round territorial aggression in a wild teleost fish"

**Legends of supplementary figures**:

**Supp. Fig. 1**: Sex differences in circulating steroid hormones during the breeding season. Dotted lines indicate the detection limit.

**Supp. Fig. 2:** Quantification of gene expression in the social brain network of breeding males and breeding females. During the breeding season, *esr1* expression was significantly higher in females in the Vs. Gene expression is shown as fold change relative to breeding males (reference group). POA: preoptic area; Vv: ventral nucleus of the ventral telencephalon; Vs: supracommissural nucleus of the ventral telencephalon; *cyp19a1b*: brain aromatase gene; *esr1*: estrogen receptor 1 gene; *esr2b*: estrogen receptor 2b gene; *ar2*: androgen receptor 2 gene.

**Supp. Fig. 3**: Sex differences in circulating steroid hormones during the non-breeding season. Dotted lines indicate the detection limit.

**Supp. Fig. 4:** Quantification of gene expression in the social brain network of non-breeding males and females. There was no significant difference in any gene in any area. Gene expression is shown as fold change relative to non-breeding males (reference group). POA: preoptic area; Vv: ventral nucleus of the ventral telencephalon; Vs: supracommissural nucleus of the ventral telencephalon; *cyp19a1b*: brain aromatase gene; *esr1*: estrogen receptor 1 gene; *esr2b*: estrogen receptor 2b gene; *ar2*: androgen receptor 2 gene.

**Supp. Fig. 5**: Correlation matrices for the four experimental groups. Spearman test. Cortisol; E1: estrone; E2: 17β-estradiol; 11-KT: 11-ketotestosterone; T: testosterone; A4: androstenedione; *ar2*: androgen receptor 2 gene; *esr2b*: estrogen receptor 2b gene; *esr1*: estrogen receptor 1 gene; *cyp19a1b*: aromatase b gene.

**Supplementary Figure 1.**

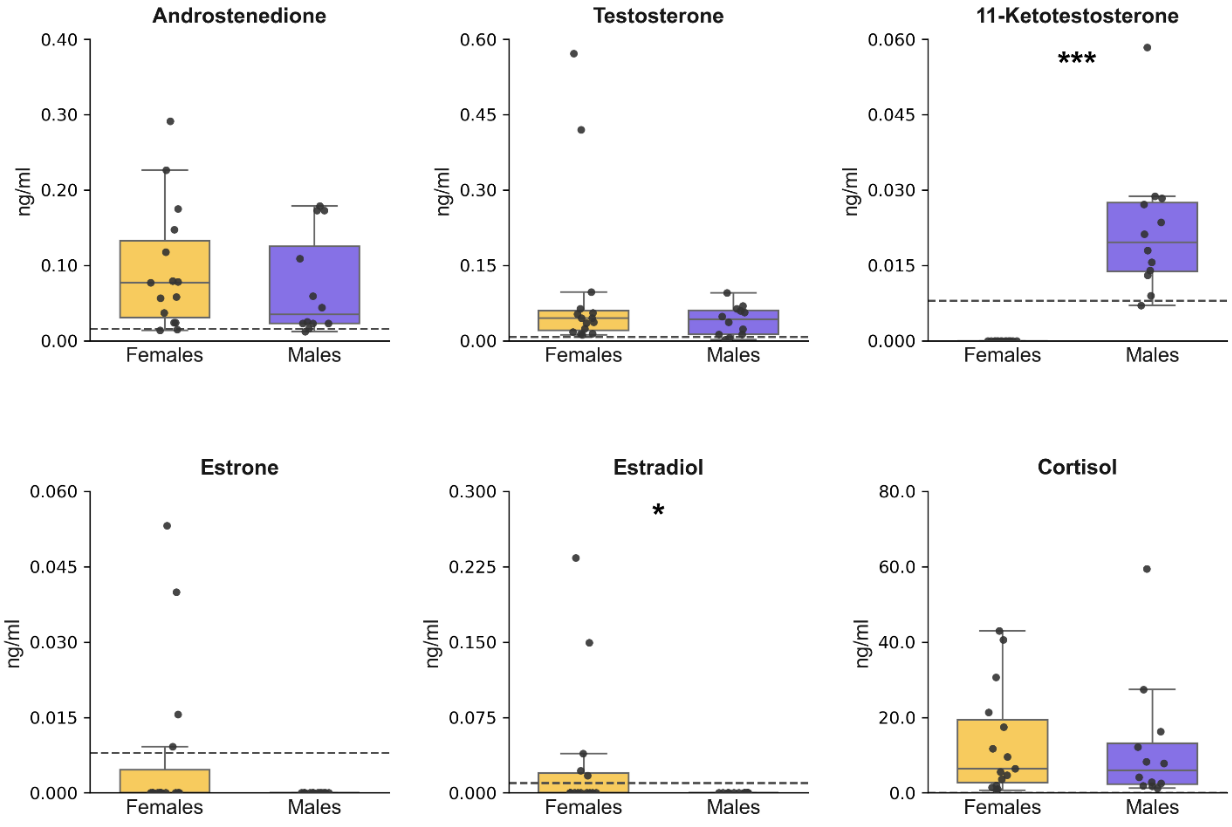

**Supplementary Figure 2.**

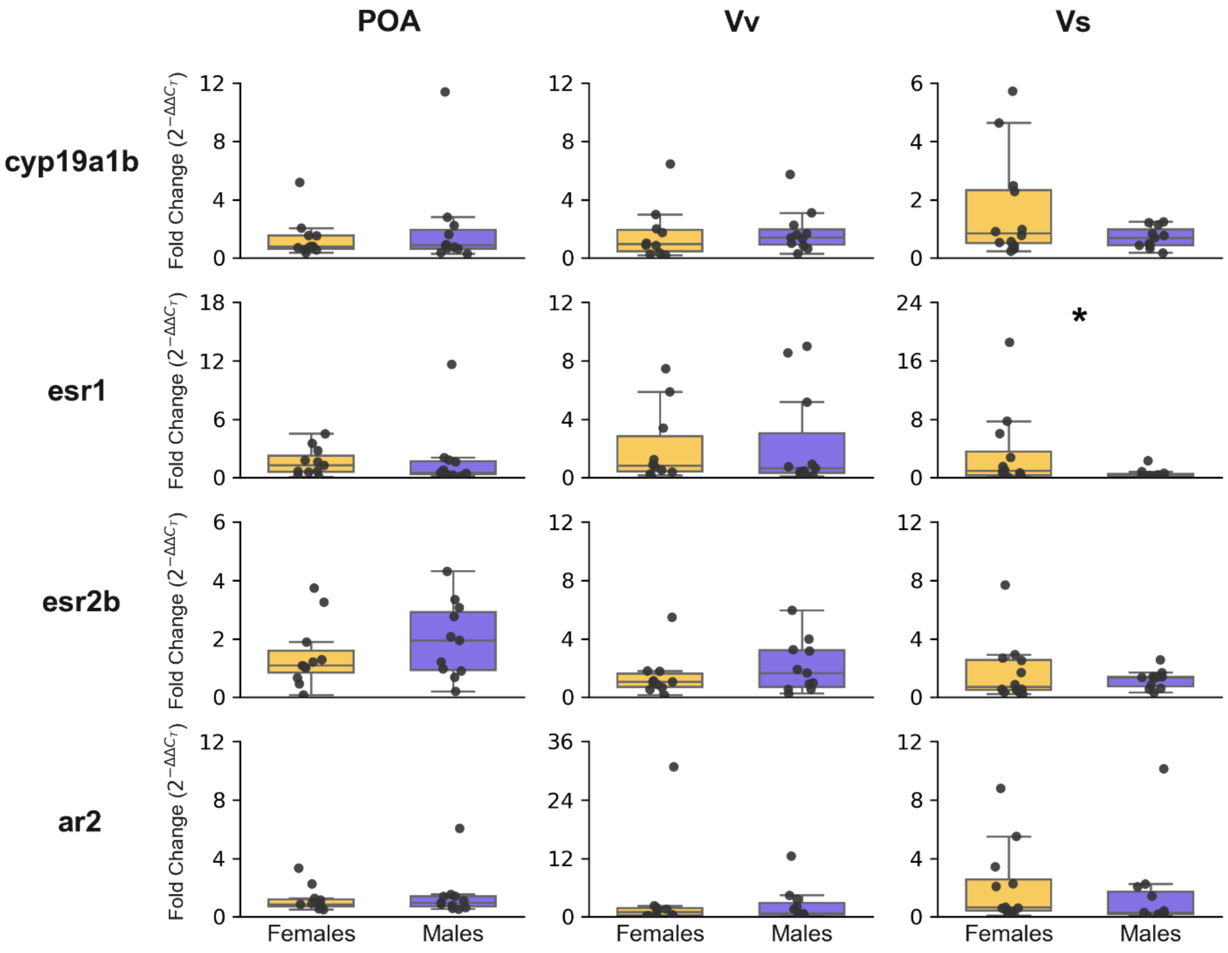

**Supplementary Figure 3.**

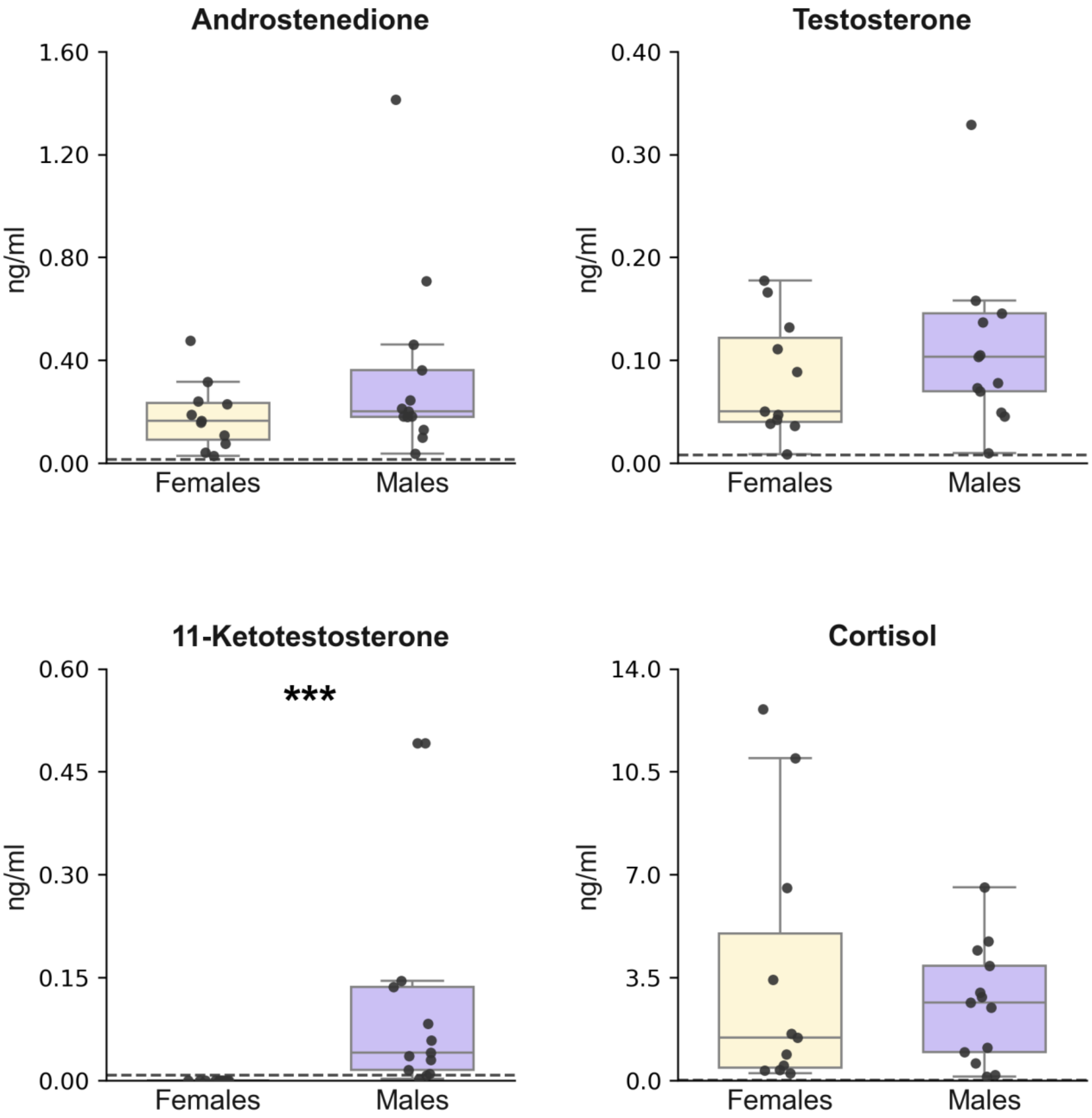

**Supplementary Figure 4.**

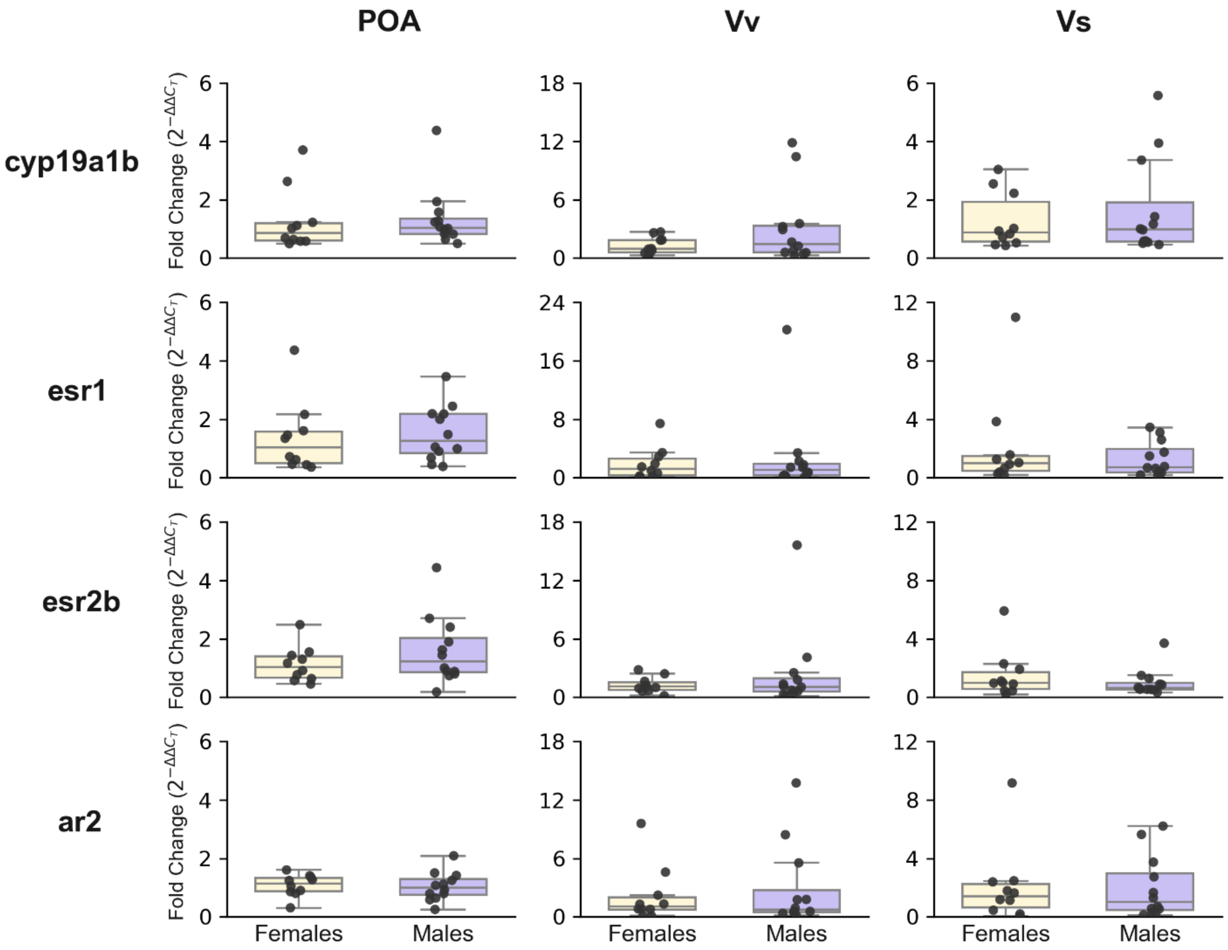

### Supplementary Figure 5.

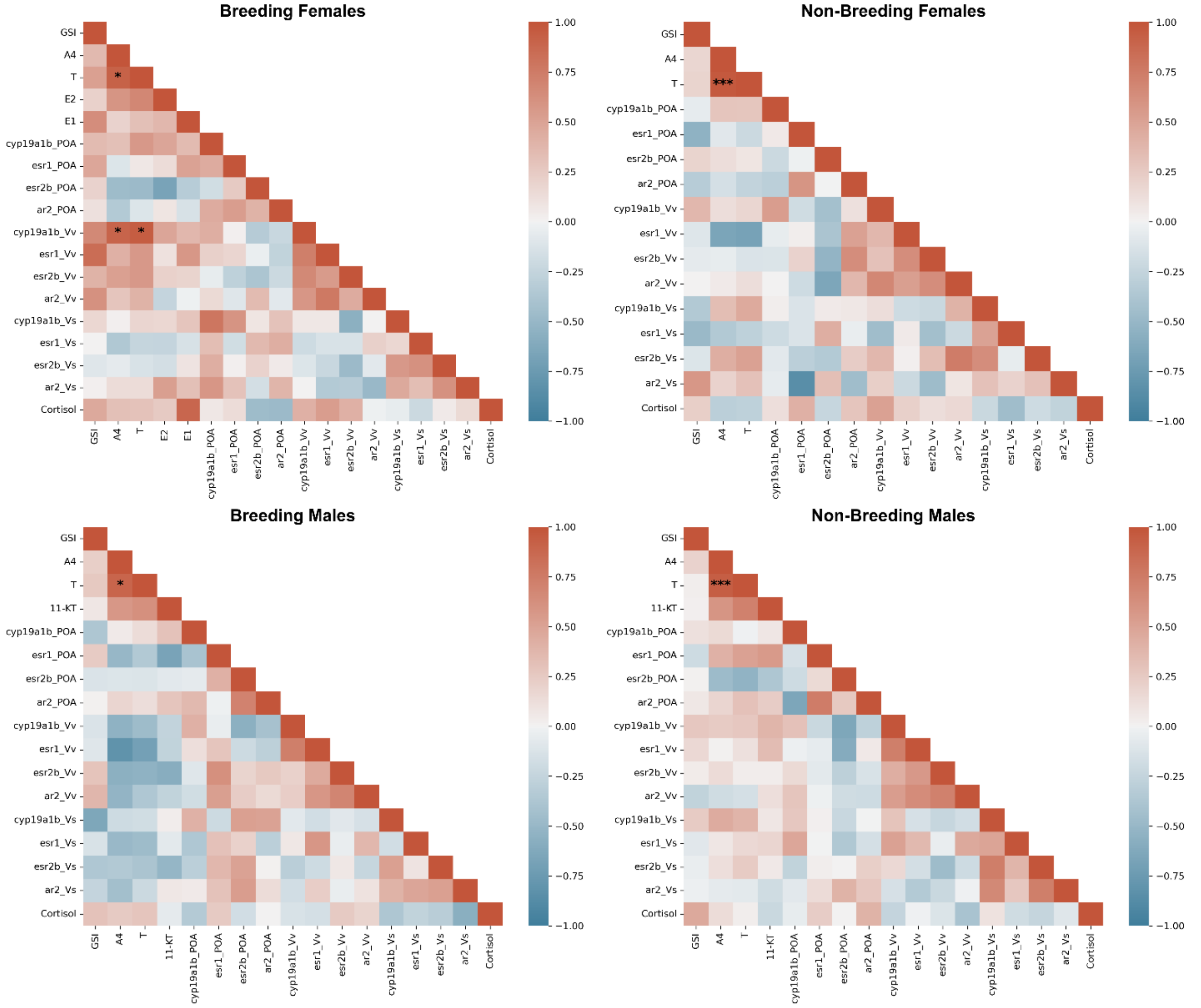
